# Self-assembling nanoparticles to assess multivalent interactions between influenza A virus hemagglutinin and glycan surfaces

**DOI:** 10.64898/2026.06.24.734152

**Authors:** María Ríos Carrasco, Daniel Y.E.L. Tambuwun, Zoé Ducarne, Hannah L. Turner, Elif Uslu, Andrew B. Ward, Geert-Jan Boons, Jurriaan Huskens, Robert P. de Vries

## Abstract

The multivalent display of surface glycoprotein hemagglutinin (HA) on Influenza A viruses (IAVs) enhances the overall binding avidity to sialylated glycans on host cell surfaces. While precomplexing HA trimers with antibodies increases multivalency and avidity, this method does not replicate the virion’s geometry and limits insights into the multivalent binding process. Here, we use perfectly controllable icosahedral protein nanoparticles to examine the multivalent HA receptor-binding properties. We compare three HA presentation systems with varying degrees of multivalency: single HA trimers, antibody-precomplexed HA trimers, and HA trimers on nanoparticles. Our results indicate that increasing HA valency enhances binding avidity across various glycan surfaces, including erythrocytes, cells, and lipid bilayers with varying glycan densities, while maintaining receptor specificity. By combining functional and non-functional HA trimers during nanoparticle formation, we create statistical mixtures of nanoparticles with varying valencies. At high receptor densities, nanoparticles with few functional trimers still bind strongly, whereas at low receptor densities, a patch of five HA trimers appears necessary for binding. As a key finding, we observe that such a statistical mixture of nanoparticles with functional and nonfunctional HAs binds to glycan surfaces in a stronger density-dependent manner than fully functional particles. We also observe differences in binding modes that correlate with the number of functional trimers, the glycan structure (linear vs branched), and the densities achievable with these glycans. Overall, our findings demonstrate that the presentation of multivalent HA plays an enormous role in the response to glycan receptor type and density, with implications for the future design of virus monitoring, viral inhibitors, and targeting vectors.

## Introduction

Each year, Influenza A viruses (IAVs) cause a high number of infections worldwide, leading to seasonal epidemics and occasional pandemics ^1^. Annual vaccines are needed to counteract the virus’ rapid mutational rate and host immune response evasion ^2^. The genetic variability of this virus is largely driven by changes in its surface proteins, hemagglutinin (HA) and neuraminidase (NA) ^3^. HA initiates viral entry by binding to sialic acid (Sia) receptors on the host cell surface, while NA disrupts these interactions to release new virions ^4, 5^.

The structure of the HA protein underlies its receptor-binding specificity, which is essential for host recognition and viral entry. HA forms homotrimers that contain head and stalk domains ^6^. The receptor-binding site (RBS) of HA is in its head domain and binds Sia ^7^. Human IAVs preferentially bind to α-2,6-linked Sia, abundantly found in the human upper respiratory tract, while avian viruses favour binding to α-2,3-linked Sia, found in the avian upper respiratory tract and human lower respiratory tract ^8^. These linkage differences in binding specificity to different numbers of galactose-N-acetylglucosamine (LacNAc) repeats and their distribution determine IAV receptor-binding specificity ^7^.

A single HA trimer binds weakly to sialylated glycans ^5^. This low affinity is overcome by multivalent interactions, which, overall, provide stronger avidity ^9^. To study the importance of multivalency in IAV binding, a commonly used approach is to precomplex HA trimers with antibodies to increase HA valency. This precomplexing method shows differential binding between avian and human HA ^10^. For instance, avian HAs required antibody precomplexation to bind to glycans. In contrast, human HAs, which can already bind elongated, multiple-LacNAc-containing glycans as single trimers, displayed broader glycan binding when precomplexed, thereby gaining the ability to bind linear glycans as well ^10^. While antibody precomplexing illustrates multiple HAs as determinants of multivalency, other structural factors can also be crucial. These factors include receptor length, receptor density, and HA-Sia orientation ^11^. These parameters have been studied by different techniques, including biolayer interferometry (BLI), glycan microarrays, and surface gradients in supported lipid bilayers, among others ^12^.

Experimental IAV vaccine design research harnesses the multivalency properties inherent to IAV ^13, 14^ by using nanoparticles (NPs) that co-display different HAs, resembling viral capsids ^15, 16^. The literature reports that the B-cell response elicited by these nanoparticle-based vaccines is broader than that of conventional influenza vaccines ^13, 14^. It has been suggested that multivalent antigen presentation strengthens interactions with B-cell receptors in HIV infection, leading to increased numbers of antigen-specific B cells ^17^. In the case of HAs, the antigen-specific antibody response to HA is greater than that to nanoparticle scaffold antibodies ^14^. Computationally designed NPs have been a fascinating addition to the field of immunogen design ^14–16, 18^ and have been successfully used for immunization against HIV and SARS-CoV-2 ^19–21^.

In this study, we examined whether these qualities could translate into the ability to study HA-glycan interactions, in particular the effect of their multivalent display on the binding avidity and glycan selectivity. We used the I53-50 NP system, which is a computationally designed two-component icosahedron with a total of 120 subunits: 60 subunits (I53-50A) assemble into 20 trimers, while the other 60 (I53-50B) assemble into 12 pentamers (Fig. 1A) ^15, 16^. Modifications to the I53-50A N-terminal enable linkage to the C-terminal of a desired antigen. These modifications make I53-50A the antigen-bearing component of an antigen-presenting NP, thereby allowing the presentation of various antigens, in our case, the HA of interest ^18^.

**Figure 1.**
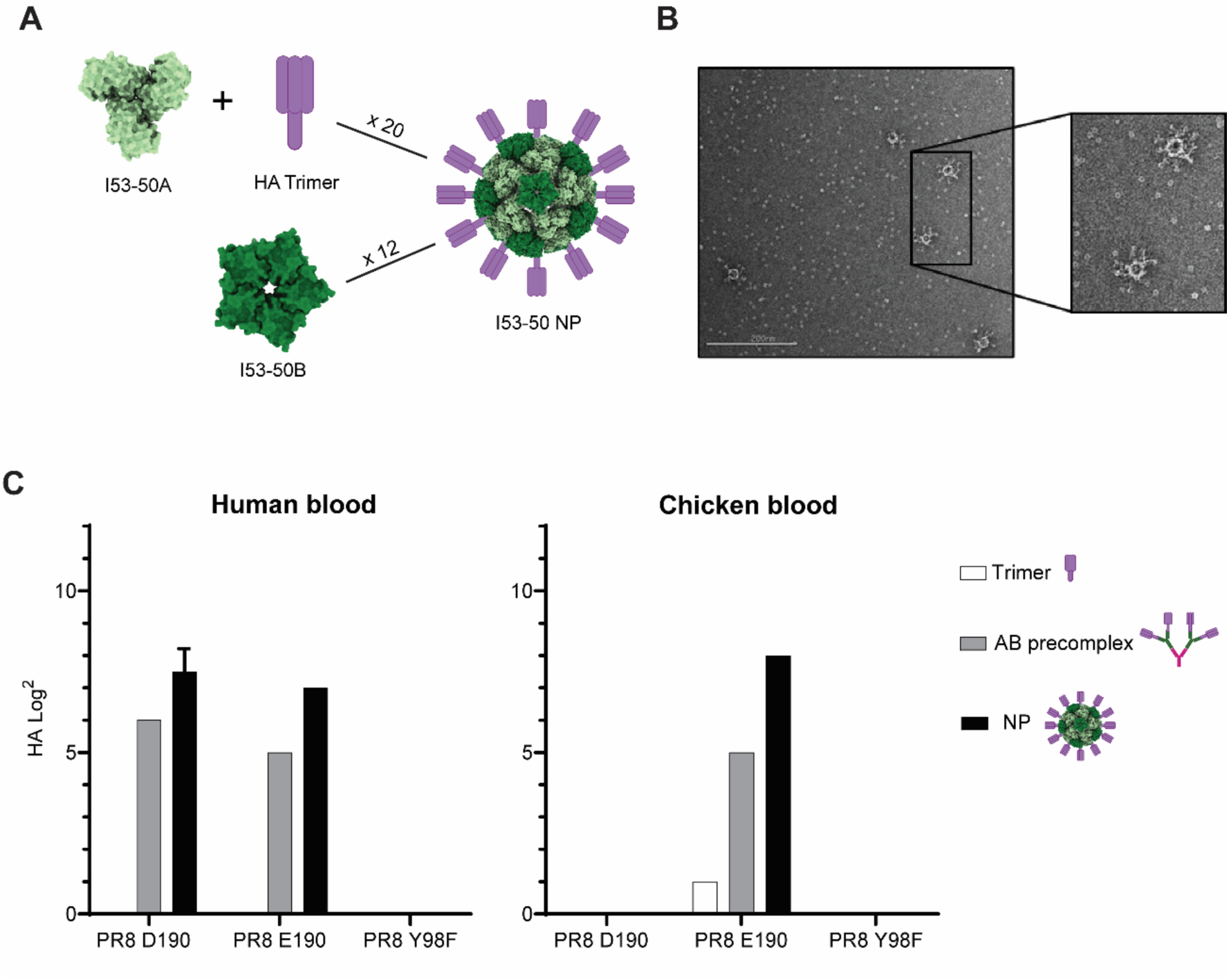
(A) Schematic representation of I53-50 nanoparticle system assembly decorated with hemagglutinin trimers. A trimeric I53-50A component is fused to the HA trimer of interest. Twenty copies of these I53-50A-HA self-assemble into twelve I53-50B pentameric components to form a complete nanoparticle. (B) nsEM micrograph of I53-50 NP decorated with PR8 D190 HA trimers. (C) Protein functionality was assessed by hemagglutination assay with human and chicken erythrocytes, comparing single trimers (white bars), antibody-precomplexed trimers (grey bars), and nanoparticle-displayed trimers (black bars).

We hypothesized that these NPs could serve as a tool to elucidate how viral protein presentation affects the multivalent binding of IAV HA to sialic acids. Modifying these NPs with functional and non-functional trimers yielded distinct binding behaviors. This suggests that NPs can serve as a tunable platform for assessing the contributions of individual HAs, depending on how interactions are arranged in space and on the constraints imposed by glycan architecture. With this work, we aim to provide insights into how HA valency and receptor density together affect binding. Our results contribute to a better understanding of how IAVs engage host cells during viral entry and highlight the importance of multivalent HA display for efficient, diverse receptor engagement.

## Results

### Nanoparticles as multivalent systems to study hemagglutinin receptor binding properties

Antibody pre-complexation of hemagglutinin (HA) increases valency relative to single trimers and is widely used in the field to boost signal in various biological assays ^10, 22^. However, it is not representative of the number and orientation of HA displayed on a virion, since neither the number nor the spherical display of HA sites can be controlled. We turned to the versatility of the self-assembling two-component I53-50 nanoparticle (NP) system ^15, 16, 20^, which enables easy fusion of our hemagglutinin of interest to the I53-50A trimeric component ^19, 20^. This allowed us to control both valency and spherical display and to access combinations of functional and nonfunctional HAs within the same construct.

Based on previous protein work ^23^, we designed three recombinant hemagglutinin constructs, each containing a SpyTag and an I5350A component, and fused an N-terminal mOrange2 molecule to each. It has been previously described that adding fluorophores N- or C-terminal to a protein can improve protein folding and stability ^24^. First, we inserted the DNA fragment encoding the hemagglutinin of A/Puerto Rico/8/1934(H1N1) strain (PR8), known to be an α2,6-linked Sia binder, into our construct. To this HA, we introduced a point mutation at amino acid position 190 (H3 numbering) from D to E, which changes specificity from α2,6-linked to α2,3-linked sialosides, leading to PR8 E190 ^25^. Lastly, we introduced another point mutation into the original PR8 at amino acid position 98, from Y to F (PR8 Y98F), known to abolish receptor binding ^26^.

The introduction of I5350A enabled presentation of these recombinant proteins on an NP using the complementary I5350B core (Fig. 1A). NP assembly was visualized by negative stain electron microscopy (nsEM) (Fig. 1B). Averaged pictures corresponded to the expected computational design originally described by Bale et al. ^16^ and resembled other immunogen design studies ^27^. We used our proteins as single trimers, precomplexed with primary and secondary antibodies, and assembled into NPs. We chose the hemagglutination assay as a standard 3D assay to assess the binding of our multivalent constructs. We employed human and chicken erythrocytes, which are abundant in α2,6- and α2,3-linked sialosides, respectively. Single trimers could not agglutinate erythrocytes, given the low avidity of single protein-glycan interactions. When precomplexed, we observed binding to the preferred receptor, α2,6-linked for PR8D190 (human blood) and α2,3-linked for PR8E190 (chicken blood). NP presentation boosted PR8 binding to human red blood cells, whereas the NP signal was significantly increased in both human and chicken blood for PR8E. As a control, PR8Y98F did not bind to any blood type, either in precomplexed or in nanoparticle form. Conclusively, the NP-HA presentation further increased binding avidity compared to antibody precomplexation.

### Surface receptor density plays a key role in different multivalent hemagglutinin presentations

To gain further insight into the constructs’ capacity to bind specific glycan structures, we performed a range of binding assays on cell lines with varying surface glycan compositions. These cell lines are Madin-Darby canine kidney (MDCK) and “humanised” MDCK (hCK). On the one hand, MDCK cells display a mix of α2,6-linked Sia on short N-glycans (one or two LacNAc repeats) and α2,3-linked Sia on a variety of N-glycans ^28^. On the other hand, hCK cells exclusively display α2,6 Sia on short N-glycans with one or two LacNAc repeats ^28, 29^.

We examined the binding properties of the PR8D190, PR8E190, and PR8Y98F-displaying multivalent systems (trimer, precomplexed, and NP) to MDCK and hCK cells using flow cytometry (Fig. 2A). Our data highlight the binding specificities of the HAs for avian- and/or human-type receptors. For this assay, we gated a viable single-cell population and measured its mean fluorescence intensity in triplicate. None of the single trimers bound either cell line. When precomplexed, PR8D190 bound more strongly to hCK than to MCDK, consistent with the presence of greater amounts of α2,6-linked Sia on its surface. There was an increase in the signal of PR8 NPs compared to precomplexed in both MDCK and hCK. However, the difference was significantly greater for MDCK than for hCK, and we confirmed that surface receptor density plays a key role in these interactions, as previously reported ^30^. Following the trend, PR8E190 NPs also bound strongly to MDCK, to a greater extent than their precomplexed counterparts. Precomplexed PR8E190 and NPs could barely bind hCK, since there is no display of α2,3-linked Sia at all. As expected, none of the PR8Y98F-displaying systems successfully bound to either cell line.

**Figure 2.**
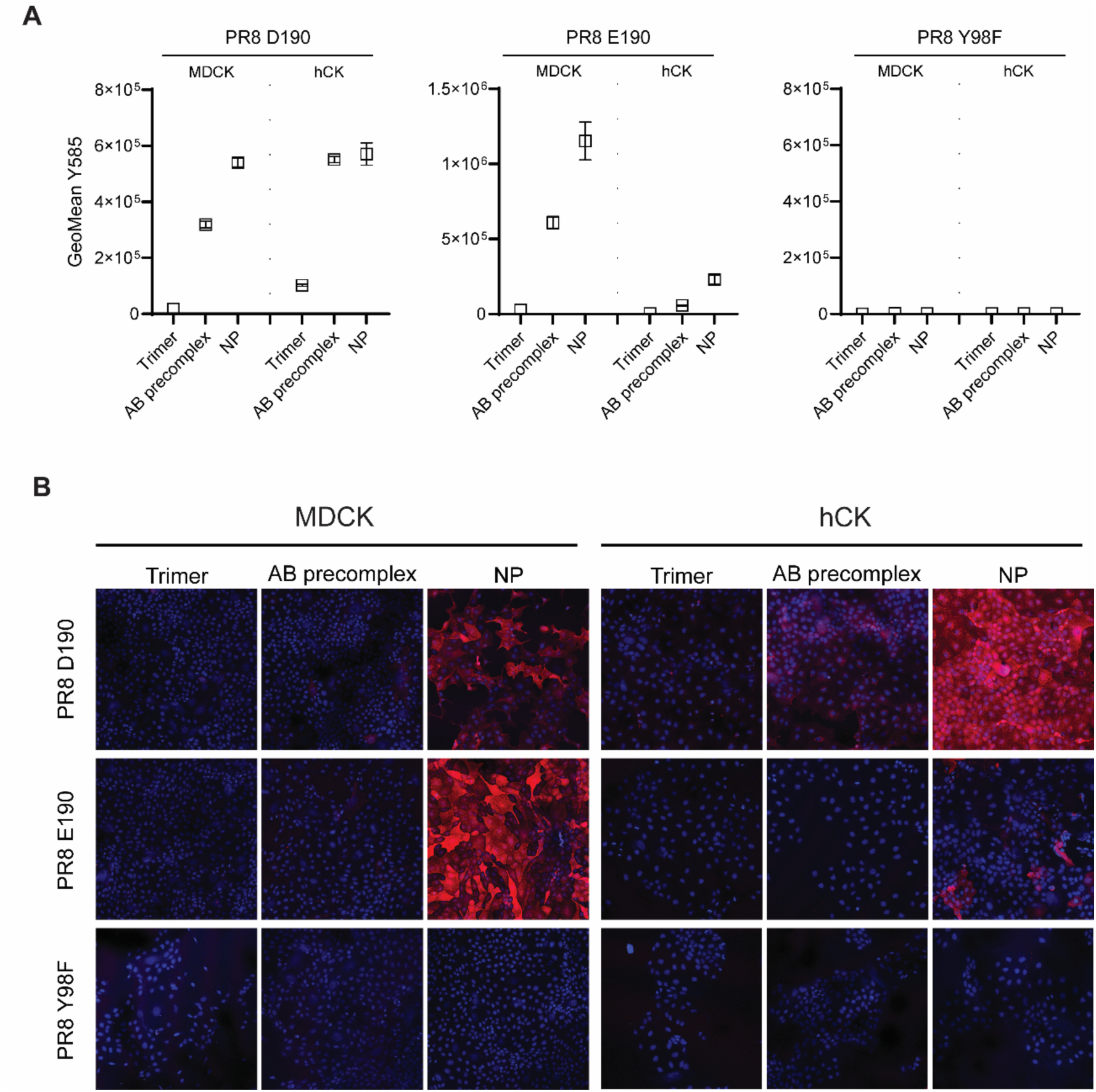
(A) Binding of PR8 D190 (binds α2,6-Sia), PR8 E190 (preferentially binds α2,3-Sia) and PR8 Y98F (does not bind sialylated glycans) using three multivalent systems (trimer, antibody-precomplexed and nanoparticle) to MDCK and hCK cells was assessed by flow cytometry. Triplicate measurements were performed; the mean fluorescence and all individual measurements are displayed. Fluorescence comes from the intrinsic mOrange2 molecule fused to each one of the I53-50A-HA trimers. (B) A comparison of these three multivalent systems was also done by immunofluorescent staining of PR8 D190, PR8 E190, and PR8 Y98F with MDCK and hCK cells.

We also performed immunofluorescent staining with the same cell lines employed for flow cytometry. Overall, higher valency resulted in increased binding from trimers to precomplexed and to NPs (Fig 2B). PR8D190 and PR8E190 NPs displayed the strongest binding to hCK and MDCK, respectively, as observed in flow cytometry. PR8 NPs could engage receptors on MDCK cells, even though these cells express only trace amounts of α2,6-linked Sia. In contrast to flow cytometry results, this assay showed that PR8E NPs bound to hCK, albeit to a lesser extent than other NPs that do not display α2,3-linked Sia. This observation highlights the potential of these NPs to pick up very weak binding interactions that are lost due to a lack of multivalency. None of the H1PR8 Y98F-displaying systems successfully bound to either cell line, as expected for this negative control.

### PR8D190 binding on planar 2D glycan surfaces

Receptor density has been reported to be a crucial factor in multivalent interactions. We performed binding studies of PR8D190 on glycan surfaces with control over receptor density. We used supported lipid bilayers (SLBs) to which streptavidin (SAv) was attached, enabling surface functionalization with biotinylated human or avian glycan receptors at controlled surface densities (Fig. 3A) ^31^. Stepwise layer formation and binding studies were monitored by QCM-D as depicted in Fig. S1. First, unilamellar vesicles (100 nm) composed of 1,2-dioleoyl-sn-glycero-3-phosphocholine (DOPC) and a pre-defined fraction of 1,2-dioleoyl-sn-glycero-3-phosphoethanolamine-N-(biotinyl) (DOPE-biotin) were used to form SLBs through vesicle fusion (step 1). Subsequently, streptavidin (SAv) was immobilized on the surface by strong biotin–SAv interaction (step 2). The surface-bound SAv retains additional unoccupied binding pockets, which were subsequently utilized for the attachment of the glycans of interest, in this case linear triLacNac α2,6-linked Sia (L-6SLN_3_, step 3). The interactions of the PR8D HA NP with glycan is specific to the α2,6-linked Sia as shown by controls (Fig. S2).

**Figure 3.**
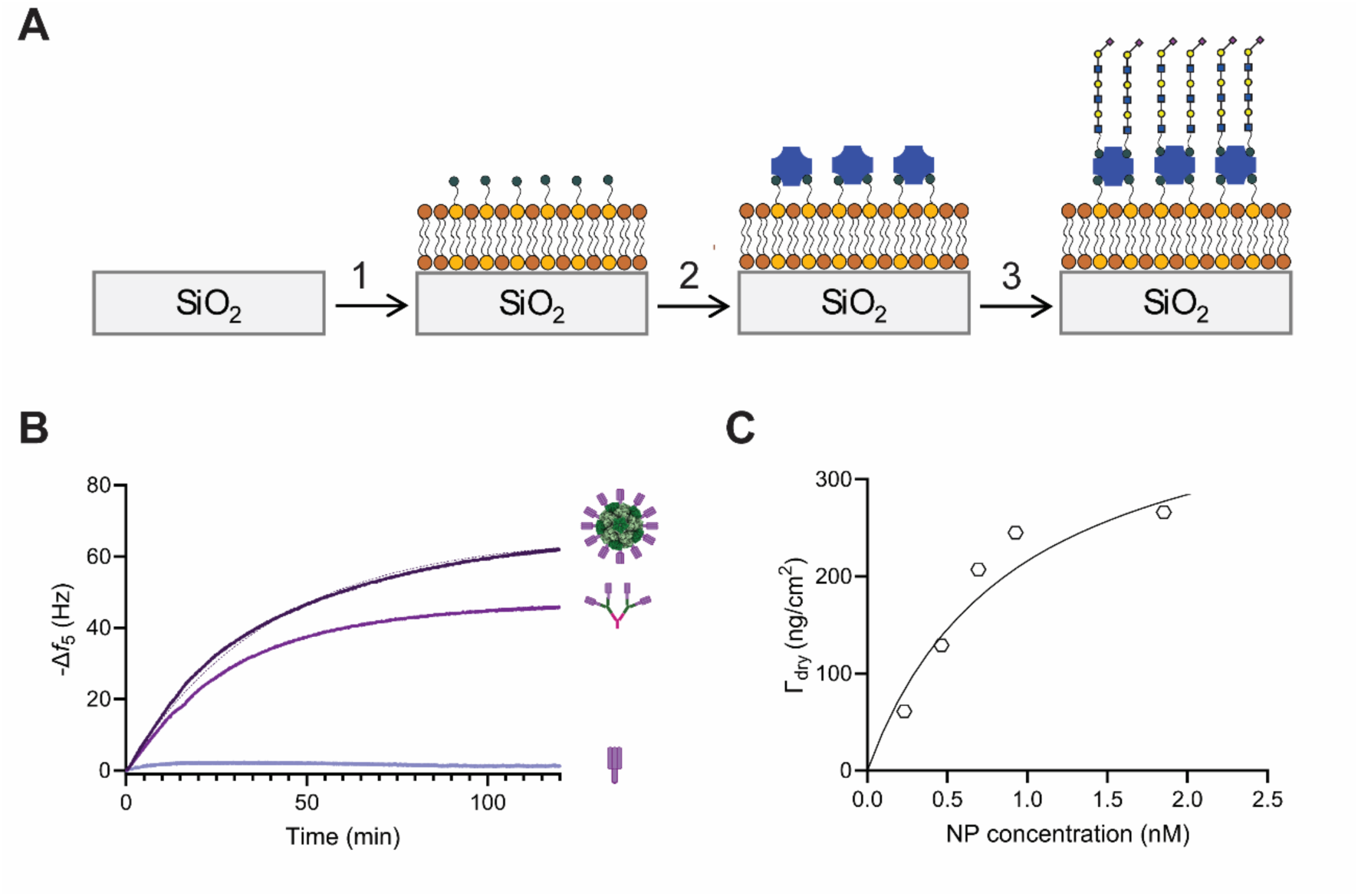
(A) Stepwise formation of (1) SLBs by vesicle fusion, (2) SAv attachment, (3) biotinylated glycan to the remaining binding pocket of SAv, (B) QCM-D traces comparison of 10 µg/mL PR8D trimer binding to linear α2,6-Sia(LN)_3_ against the same concentration of the PR8D190 in its multivalent display (*i*.*e*. in antibody-precomplexed and NP forms). The equilibrium -Δ*f*_5_ values are obtained after fitting with single exponential. (C) Dry mass of adsorbed PR8D190 NPs as a function of the NP concentration in solution upon conversion of QCM wet mass data into dry mass using the calibration curve shown in figure S3.

The binding of PR8D190 to L-6SLN_3_ was enhanced with increasing valency. The surface coverages of both antibody-precomplexed and NP were higher compared to the trimeric form of PR8D190 at 10 µg/mL on densely packed SAv surface (σ_SAv_ = 3.8 pmol/cm^2^) (Fig. 3B). Although the reported frequencies can be used to derive adsorbed masses, a quantitative comparison between multivalent forms is difficult due to differences in molecular weights and changes in hydration at higher coverages. Nevertheless, PR8D190 binds significantly only in its multivalent forms.

To better quantify surface coverage and affinity, we combined QCM with spectroscopic ellipsometry (SE) to determine dry mass coverage from the measured frequency shifts. We observe that the dry mass deviates from linearity as the frequency shift responses increase (Fig. S3a) and the hydration percentage decreases with increasing coverage (Fig. S3b). This hydration behavior can be explained by the hydrodynamic coating model, in which overlapping hydration coats of individual NPs reduce the increase in wet mass with increasing coverage ^32^. Moreover, we could also verify the layer thickness of densely-packed bound NPs on surface from the Δ*D_n_*/-Δ*f_n_* ratio (or the acoustic ratio, that is about 61 nm, see accompanying text of figure S4 for details). Therefore, we combined QCM with SE to quantify the coverage and packing of NPs on planar surfaces.

The NP assembly of PR8D190 binds to L-6SLN_3_ with an apparent affinity in the nM range. In the NP formation step, PR8D190 was added in excess (7.5-fold; see Fig. S5 for details); we therefore assume that the resulting NP concentration reflects full conversion of CompB into NPs. Dry-mass conversion of the NP coverage yields *K*_d,app_ = 1.1 nM (Fig. 3C). This apparent affinity was obtained at the maximum L-6SLN_3_ density achievable on densely packed SAv surface (σ_SAv_ = 3.8 pmol/cm^2^, σ_L-6SLN3_ = 7.6 pmol/cm^2^). We used a Langmuir isotherm to fit the data, although the true binding mechanism may deviate from it due to, for example, steric hindrance on the surface and kinetic effects. All in all, this affinity is about 6 orders of magnitude higher compared to the (mM) monovalent affinity of HA-glycan interactions (40, 41), but still lower compared to the avidity of a full virus, which is in the pM range (42).

### Variation of the number of functional trimers within nanoparticles gives insight into the 3D binding mode

In the previous sections, we showed that higher valency increases HA binding avidity, especially when HA trimers are presented on NPs. Based on this principle, we aimed to use the NP platform itself as a controllable system to create variations in HA valency – by displaying a mixture of functional and non-functional HA trimers on its surface – and studying the effects on their binding to various Sia-functionalized platforms. The functional HA trimer was PR8D190, chosen for its high specificity towards α2,6-linked Sia. The non-functional trimer was PR8Y98F, used as a negative control because it does not bind Sia.

Since the orientation of HA-Sia interactions is an important property for multivalent binding ^11^, we used two titration-based approaches (Fig. 4A). We make separate stocks of fully functional and fully nonfunctional NPs, and mix these stock solutions at different ratios, designating the resulting mixtures as pure NPs with varying fractions of functional HA. In the second approach, we designed an experimental setup in which the total amount of HA remained constant while we varied the ratio of functional to non-functional HA within a single NP batch, termed mixed NPs. We expect to obtain a statistical mixture of NPs with varying numbers of functional and nonfunctional HA trimers, corresponding to the ratio used to assemble the NPs. In all cases, the CompB concentration was kept constant, ensuring that the total NP concentrations remained constant.

**Figure 4.**
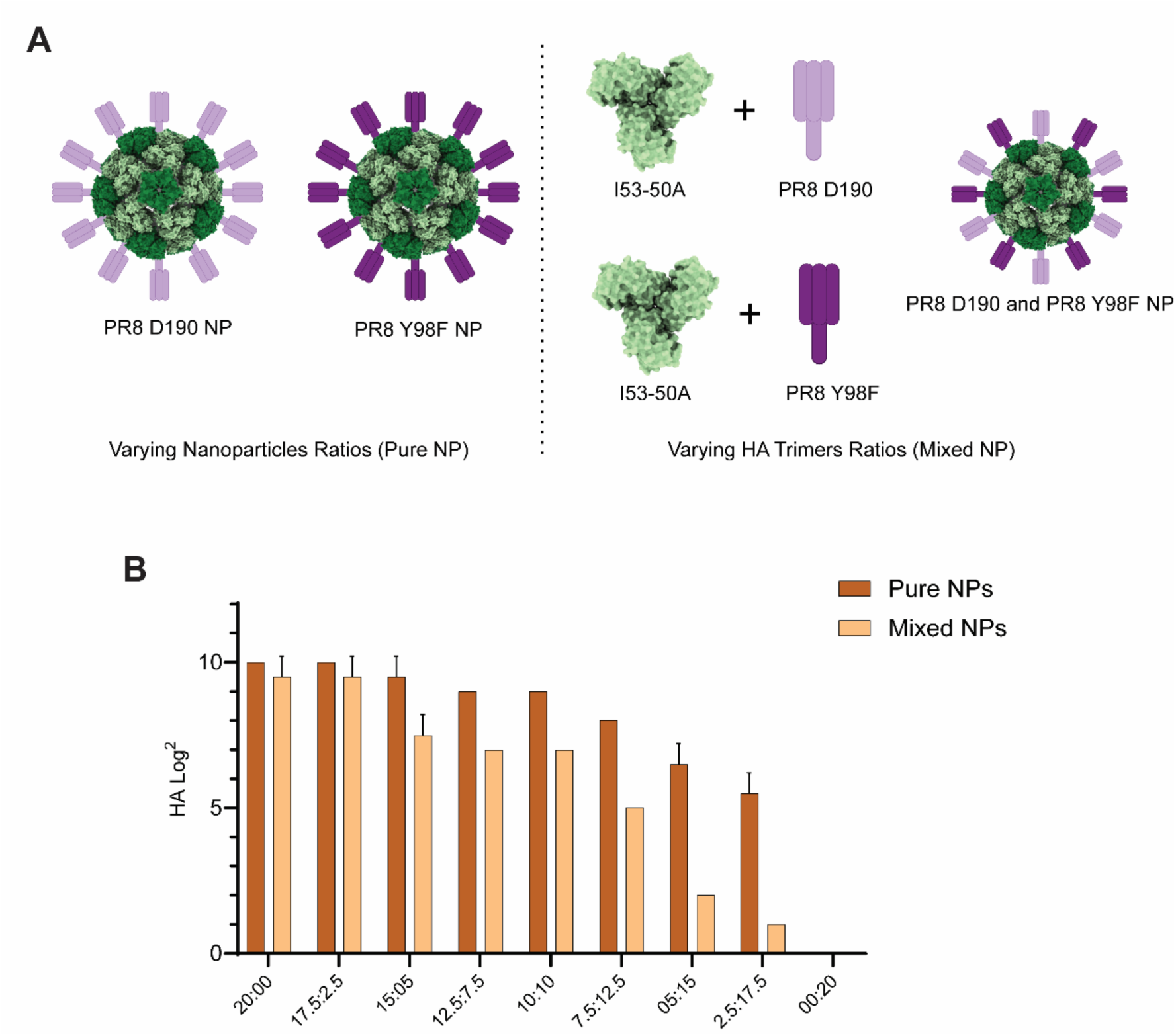
(A) Schematic representation of the NPs used for titrations. On the left, I53-50 NPs are assembled with either the PR8 D190 or the PR8 Y98F-I5350A trimeric component. Pure NPs were then mixed at varying ratios. On the right, PR8 D190 and H1PR8 Y98F-I5350A trimeric components were mixed at varying ratios prior to NPs assembly, resulting in mixed NPs. (B) Hemagglutination assay with human erythrocytes using different ratios of fully functional and fully non-functional nanoparticles (“pure NPs”) or particles assembled from a mixture of functional and non-functional HA trimers (“mixed NPs”) at a constant total NP concentration.

We performed a hemagglutination assay to study 3D NP-glycan interactions, as erythrocyte agglutination creates a mesh that allows NPs to bind on multiple sides. In the pure NPs, we observed a relatively constant binding to human erythrocytes as the amount of functional HA-trimer-displaying NPs decreased (Fig. 4B). In the mixed NPs, however, we observed a rapid decline in binding to human erythrocytes as the amount of functional HA trimer displayed on the NP decreased (Fig. 5B, 20:00 vs 15:05 ratio). As expected, binding was abrogated when only H1PR8 Y98F-displaying NPs were present in both pure and mixed NPs (00:20 ratio).

**Figure 5.**
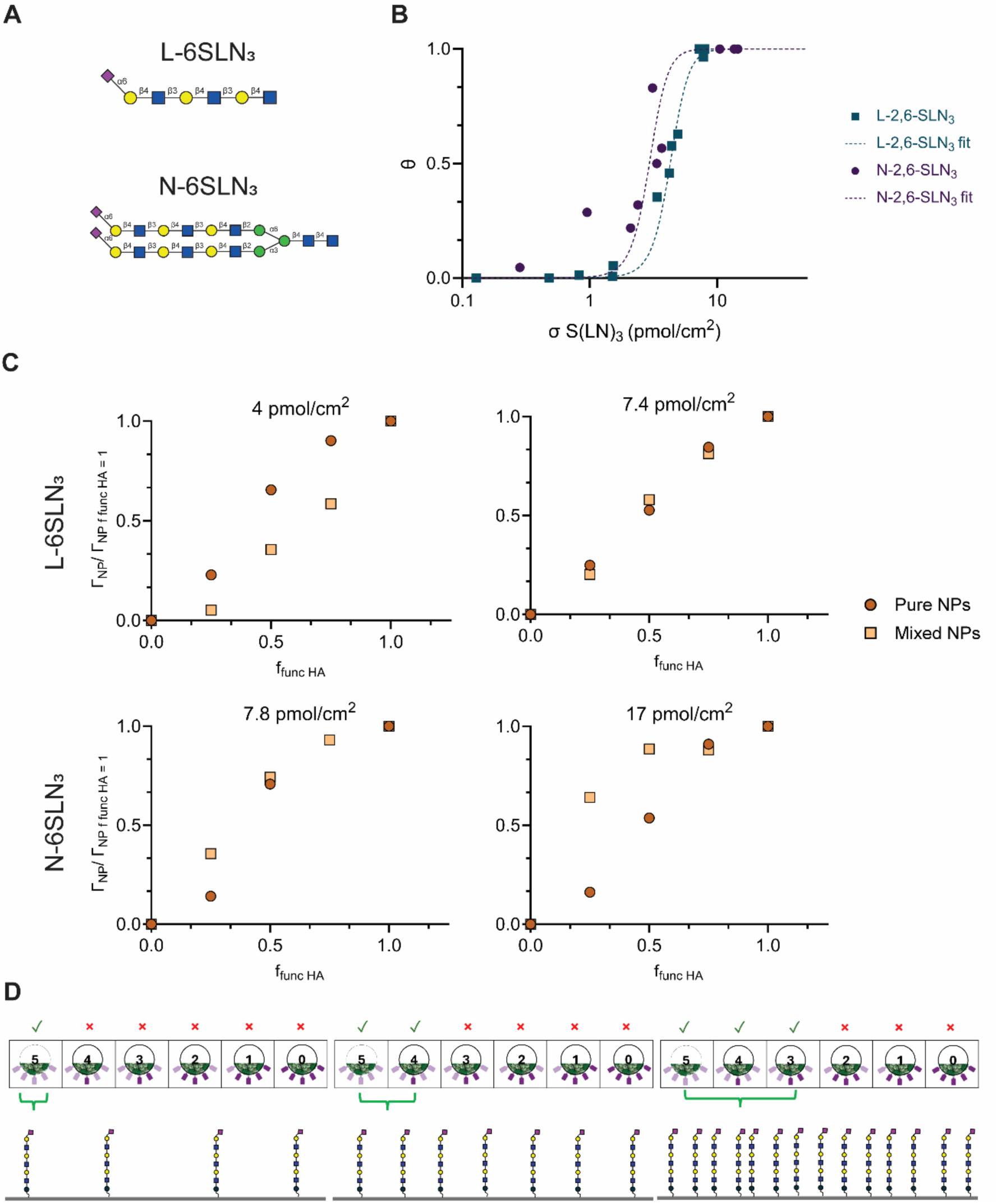
(A) Glycan structure of the linear (L-6SLN_3_) and biantennary triLacNac α2,6-linked Sia (N-6SLN_3_). (B) Titration of linear and branched glycan at varying Sia density. Density variation was achieved by varying %mol of biotinylated lipid to the lipid mixture in the vesicles. Concentration of the NP was kept constant at 1 nM. (C) Binding of pure and mixed NPs on SLBs at varying surface receptor densities of L-6SLN_3_ and N-SLN_3_ measured by QCM. The resulting - Δ*f*_5_ values were converted to dry mass coverage using the calibration curve in Figure S3. The total concentration of NP was kept constant at 1 nM while varying the fraction of functional and non-functional HA. (D) A scheme illustrating the dependency of NP binding on the product of HA and Sia densities. At low Sia density, only NP with a high patch size (*i.e.,* 5 HAs at the contact area) can bind. As the Sia density increases, particles with smaller patch sizes can bind.

These observations illustrate that, for 3D interactions, fewer NPs are needed for successful attachment in the pure NPs, since the functional trimers present work multivalently to interact with red blood cells. By contrast, when we use mixed NPs, the presence of non-functional trimers within the NPs inhibits the multivalent effect, thereby decreasing binding to erythrocytes. We hypothesize that pure NPs preserve binding functionality on all sides, allowing them to function as crosslinkers between cells, whereas mixed NPs have an increasing fraction that retains only a single available binding side, reducing their crosslinking capability. In the next sections, we address this hypothesis.

### Variation of the number of functional trimers within nanoparticles reveal differential 2D binding to linear and branched glycans

To elucidate 2D binding, we first investigated the binding of fully functional NPs to SLBs with controlled glycan densities. We employed linear triLacNac α2,6-linked Sia (L-6SLN_3_) and biantennary triLacNac α2,6-linked Sia (N-6SLN_3_), depicted in Fig. 5A. The NP presentation of PR8D190 HA showed a non-linear, superselective binding response with respect to the density of sialylated glycan receptors due to multivalency (Fig. 5B). The nanoparticle concentration was kept constant at 1 nM, and the receptor density was calculated according to the Sia density, which was either twice or four times the density of SAv in the case of L-6SLN_3_ or branched N-6SLN_3_, respectively. The data points of NP coverage *vs*. Sia density were fitted using Equation S3 by assuming 15 binding HA sites available at the contact area (Fig. S7). Here, we define the threshold receptor density (σ_threshold_) as the sialic acid density required at 50% NP coverage. The σ_threshold_ of N-6SLN_3_ appears at 2.9 pmol Sia/cm^2^, which is lower than for the linear glycan counterpart (σ_threshold_ = 4.2 pmol Sia/cm^2^). Introducing branching to the glycan thus increases the probability of intramolecular binding (*i.e.,* the effective molarity increases, Table S1). However, the maximum selectivity factor 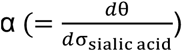 is comparable for both L-6SLN_3_ and N-6SLN_3_ (Fig. S8). This still implies that the NPs bind superselectively on both glycan types (*i.e*., there are σ_sialic acid_ ranges where α > 1 in both cases). Nevertheless, for the N-glycan, the superselective binding regime already appears at a lower Sia density, compared to the linear glycan counterpart. Thus, valencies, receptor densities, and the structural configuration of both the NP and the glycan of interest affect the enhancement of avidity.

For comparison with our previous 3D hemagglutination approach, we also studied the effects of surface receptor density on the binding of pure and mixed NP systems in a 2D setup. We observed that pure NPs bind favorably at lower Sia density, while mixed NPs are more favored at higher receptor density. We used the same SLB-SAv platform to control the Sia density. At lower Sia density (4 pmol/cm^2^), mixed NPs bind less than pure NPs at all measured ratios (Fig. 5C, top left panel). The NP coverage becomes similar as the Sia density of L-6SLN_3_ increases (Fig. 5C, top right panel). In this case, the Sia densities are limited to the maximum packing of SAv, it is not possible to reach beyond 8 pmol/cm^2^ with L-6SLN_3_. The Sia density-dependent binding is also observed with the branched N-6SLN_3_. Pure NPs and mixed NPs showed similar binding at 7.8 pmol/cm^2^ of Sia of N-6SLN_3_ (Fig. 5C, bottom left panel). This is comparable to the trend observed with L-6SLN_3_ at a similar Sia density. At the maximum density of N-6SLN_3_ (equivalent to 17 pmol/cm^2^ of Sia), mixed NPs bind more efficiently than pure NPs at all ratios measured (Fig. 5C, bottom right panel). Overall, the branching in the glycan structure simultaneously increases the Sia surface density and enables the binding of lower-valent mixed NPs.

### Mixed NPs bind more selectively than pure NPs

A more detailed analysis of the mixed NPs and their binding provides a quantitative assessment of the minimum number of HA trimers required and offers a rationale for the observed difference in binding performance between pure and mixed NPs at different glycan densities. In general, binding efficiency depends on the concentrations of both binding partners. For example, the extent of complex formation described by a 1:1 equilibrium constant is controlled by the product of the concentrations of both binding partners. Here, the interaction between an HA NP and a Sia surface is similarly controlled by the product of both of their densities, as well as by the total NP concentration. As a result, the minimum number of HAs required at the contact area between an HA NP and the Sia surface to result in sufficient avidity gain depends on the Sia surface density. As explained in the SI, we assume that our icosahedral NPs interact through a binding patch containing 5 HA trimers (Fig. S7). For NPs with a varying functional HA fraction, we derived distributions of the fractions of these mixed NPs that contain at least one patch with a defined minimum number of functional HA trimers (an extensive explanation is provided in the SI). As shown in Fig. S9, these distributions start at 100% at high functional HA fractions but decrease in a nonlinear, sigmoidal fashion as the functional HA fraction is reduced. Notably, the midpoint at which half of the mixed NPs have a patch with a given number of HAs decreases from 60% functional HA for a patch with 5 functional HAs to <5% functional HA for a patch with 1 functional HA. At comparably low Sia density, the NPs thus require a relatively high number of functional HA trimers at the binding patch to bind. In the case of mixed NPs, the fraction of particles with that required number of HAs only appears at relatively high fractions of functional HA. For pure NPs, instead, the fraction of sufficiently strong-binding NPs – which is the fraction of functional NPs, which all have a binding patch of 5 functional HA trimers – is determined directly by the functional HA fraction, which is a linear dependence.

We can now use a comparison between the binding of the pure and mixed NPs at varying Sia densities (data of Fig. 5C) to derive quantitative information about the minimal binding patch numbers for the mixed NPs. Following the analysis in the SI, we can derive the relative concentrations of particles in the mixed NP samples that bind, as a function of both the HA fraction and the Sia density (Fig. S10). When comparing these binding NP fractions with the expected functional fractions at varying minimal binding patches (Fig. S9), we see that a patch number of 5 (*i.e*., 5 functional HA trimers are needed to be located at the same side of a particle to make it bind) is required at the lowest Sia density, while this number goes down to 3 for the highest Sia density used here. Consequently, for the mixed NPs, the fraction of particles with several HAs at the binding patch equal to or greater than the required minimum remains high until relatively low fractions of functional HA. In contrast, the fraction of active NPs in the pure NPs drops more quickly, leading to a lower response. Overall, this model explains the reversal of trends in binding of pure vs mixed NPs across varying Sia densities and provides quantitative information about the necessary HA trimer densities as summarized in Fig. 5D.

A very important lesson that can now be drawn from these observations: mixed NPs are more superselective than pure NPs. This is confirmed by plotting the binding of the mixed NPs at varying Sia densities and as a function of the functional HA fraction in a superselectivity plot, like Fig. 5B (see Fig. S11). This figure shows that the superselectivity parameter α is higher at lower HA fractions. Since the superselectivity describes the sensitivity of binding with respect to varying Sia densities, this shows that mixed NPs respond more strongly than pure ones to variations in the receptor density. Qualitatively, this is evident from the trend shown in Fig. 5C and discussed above: in comparison to pure NPs, mixed NPs bind better at higher Sia density and worse at lower Sia density. In essence, this behavior is determined by the necessary HA site density to bind to a surface with a particular Sia density: lower Sia densities require higher patch numbers, and in the mixed NPs, the fraction of particles that have such a minimal patch size decreases nonlinearly with the functional HA fraction. This may provide a new paradigm for multivalent drug targeting: when targeting, for example, cells with overexpressed receptor densities, one may achieve better contrast in binding to diseased cells relative to healthy ones by using targeting vectors with randomly distributed ligands rather than a constant ligand density.

Additionally, the model can provide more information about 3D binding. Hemagglutination is such a 3D experiment, where the readout is induced by NPs crosslinking cells. Hence, binding of an NP is not enough; after binding to one cell, there still needs to be a binding patch available at the other side of the NP to bind to another cell. In the model, this requirement for two binding patches has also been evaluated (Fig. S9b). Obviously, compared to 2D binding, 3D binding requires a higher fraction of functional HA to reach the same fraction of particles capable of binding. The rest of the analysis proceeds similarly to that described in 2D. Qualitatively, the hemagglutination data shown in Fig. 4B show more efficient binding of the pure NPs, indicating that the Sia density on these cells is relatively low, requiring a high HA site density on the NPs. More quantitatively, this behavior suggests the need for 5 functional HA trimers in the contact area.

## Discussion

In this work, we proposed using nanoparticles as tunable tools to study HA-glycan interactions. We compared three systems with varying degrees of multivalency, including single HA trimers, HA trimers precomplexed with antibodies, and HA trimers displayed on nanoparticles. We aimed to investigate how multivalency affects HA-binding avidity and its relationship to receptor density. We reported that increasing HA valency enhances binding to diverse glycan surfaces while maintaining receptor specificity. By varying the ratio of functional trimers on nanoparticles, we showed that receptor density modulates the HA binding mode.

The comparison between antibody precomplexing and nanoparticle display confirmed that increasing the number of trimers increased multivalency, which boosted binding avidity, as previously observed ^10^. This observation aligns with the binding trends of HA-presenting rosettes and whole IAV viruses to synthetic glycan surfaces ^33^. In line with previous studies, we report here that glycan architecture influences HA’s multivalent binding. Symmetrical N-glycans have a structure that allows for bidentate binding, while linear glycans do not ^10, 34^. Additionally, in a recent study, Liang et al. show that human-type HAs bind symmetric and asymmetric N-glycans similarly, suggesting other glycan structures can engage in bidentate binding ^35^.

A potential limitation of the I53-50 nanoparticle system is that it is smaller than an actual influenza A virion and has trimers evenly assembled on its surface, which does not correspond to HA clusters reported for IAV ^36, 37^. These clusters are known to contribute to HA multivalent interactions with host cell receptors ^12, 37^. Therefore, aside from confirming the NP assembly with nsEM, we cannot directly relate the effects of both the orientation and distribution of the displayed HA trimers on the nanoparticle surface to those observed in actual viruses. This orientation and clustering will likely affect glycan binding, as they determine how accessible the RBS of each trimer is to different glycan species ^12^. Nevertheless, the I5350A nanoparticle offers a well-defined composition and, as shown in this and other studies, enables the combination of trimers with multiple specificities in varying ratios ^38^. These advantages demonstrate the system’s potential for HA receptor-binding studies in a controlled setting, which is particularly relevant considering recent findings that influenza A viruses actively remodel the nanoscale organization of the host cell surface during attachment ^39^. Future research approaches can focus on displaying multiple HA subtypes to assess multivalency across IAV strains within a single system.

In addition to demonstrating the new application of nanoparticle systems for receptor-binding studies, our findings show the high versatility of the I53-50 nanoparticle system. Our version of the I53-50A component contains a SpyTag, part of the SpyTag-SpyCatcher system ^40^. These constructs are thus compatible with other multivalent systems that could complement our observations by varying the number of HA trimers displayed ^41^. On the receptor side, lipid bilayers have been an excellent tool for precise control of receptor density ^33^. Expanding on glycan densities and species across different ratios can be an insightful approach for further studying HA-glycan interactions.

A unique feature of the NPs is the ability to control the (average) HA site density, which cannot be achieved with full viruses. Here, we show that both HA site density and Sia density control binding efficiency in both 2D surface binding and 3D hemagglutination. Our findings indicate that only a few trimers are sufficient for effective attachment of nanoparticles to host cells. However, by altering both the number of functional trimers on nanoparticles and the surface receptor density, we revealed distinct multivalent binding regimes across Sia densities, which can be achieved with linear and branched compounds. The overall receptor density appears to be the unifying factor explaining differences in binding between pure and mixed NPs. Most intriguingly, mixed NPs can be more selective to receptor density than pure NPs, which may hold promise for future work, for example, in the design of targeting vectors. We believe that multidisciplinary approaches, such as this study, which combines surface chemistry, virology, and glycobiology, are essential for a better understanding of IAV binding and for expanding the toolbox of techniques for receptor-binding studies.

## Methods

### Expression and purification of hemagglutinin

Codon-optimized cDNAs encoding hemagglutinin (HA) from distinct HA strains were cloned into the pCD5 expression vector as previously reported ^23^. The pCD5 vector was engineered to allow insertion of HA-encoding cDNAs in-frame with sequences encoding an N-terminal secretion signal, the fluorescent protein mOrange2 positioned at the N-terminus, a GCN4 trimerization motif (RMKQIEDKIEEIESKQKKIENEIARIKK), and a Twin-Strep tag (WSHPQFEKGGGSGGGSWSHPQFEK; IBA, Germany) ^23, 42^.

Additional modifications to the pCD5 backbone included the incorporation of a SpyTag ^40^ and the I5350A trimeric component, which is needed for nanoparticle assembly ^16^. The resulting constructs were transfected into HEK293S GnTI(−) cells, and expressed proteins were purified as previously described ^23^. Briefly, plasmid DNA and polyethyleneimine I (PEI) was combined at a 1:8 (µg DNA:µg PEI) mass ratio and added to the cells. After 6 hours, the transfection mixture was replaced with 293 SFM II suspension medium (Invitrogen, 11686029) supplemented with 2.0 g/L glucose, 3.6 g/L sodium bicarbonate, 3.0 g/L Primatone (Kerry), 1% GlutaMAX (Gibco), 1.5% DMSO, and 2 mM valproic acid. Supernatants were collected five days post-transfection, and recombinant proteins were purified using Strep-Tactin Sepharose beads (IBA Life Sciences) according to established protocols ^43^.

### Nanoparticle visualization by negative stain EM

Nanoparticles in 10mM Tris, 150mM NaCl at 4°C were added to homemade carbon-covered copper grids (EM Sciences) and stained with 2% uranyl formate. Negative stain images were collected on a 120KV Spirit microscope equipped with an Eagle 4K CCD (Thermo Fisher). Leginon (PMID 10406132) software was used for automated collection and micrographs were viewed via Appion (PMID 19263523).

### Hemagglutination assay

Hemagglutination assay was performed as described previously ^23^. HA proteins were precomplexed with α-strep-tag human antibody (GenScript) and goat-α-mouse human antibody (Novus Biologicals) for 30 minutes at a molar ratio of 4:2:1, respectively. For nanoparticle assemblies, HA proteins were pooled with CompB in PBS overnight at 4 °C. Human and chicken erythrocytes were washed twice with PBS and diluted to 1% solution. Serial dilutions of protein mixes, precomplexed and NPs, were incubated with the red blood cells for at least 1 hour at room temperature before visualizing results.

### Flow cytometry

Flow cytometry experiments were performed as previously described ^44^. Madin-Darby canine kidney (MDCK) cells and humanized MDCK (hCK) cells were washed with PBS supplemented with 3% BSA and 2 mM EDTA, and resuspended to a final concentration of 500,000 cells/ml. A total of 50.000 cells were added to 96-well U-bottom plates (Falcon). Protein mixes were added to cells, which were then incubated on ice for 90 minutes. Cells were then washed with PBS supplemented with 3% BSA and 2 mM EDTA and incubated with a 1:2000 dilution of eBioscience Fixable Viability Dye eFluor 780 (65-0865, Thermo Fisher Scientific) for 5 minutes on ice. Cells were resuspended in a final volume of 100µl, and flow cytometry was performed using CytoFlex (Beckman Coulter) using appropriate laser voltages. Data were analyzed with FlowJo using a gating strategy to select single, live cells and calculate the fluorescent geometric mean of triplicate values.

### Protein histochemistry

Histochemical analysis was performed as previously described ^45^. Paraffin-embedded tissue sections from human and chicken trachea were re-hydrated with 100%, 95%,70% ethanol and distilled water. Antigens were retrieved with warm citric acid at pH 6.0, and endogenous peroxidase was blocked with 1% H202 in methanol for 20 minutes at RT. Tissues were washed with PBS-Tween and incubated in 3% BSA overnight at 4 °C. HA proteins were precomplexed with α-strep-tag human antibody (GenScript) and goat-α-mouse human antibody (Novus Biologicals) for 30 minutes at a molar ratio of 4:2:1, respectively. For nanoparticle assemblies, HA proteins were pooled with CompB in PBS overnight at 4 °C. Protein mixes were added and incubated for at least 90 minutes at room temperature. Slides were washed with PBS, incubated with DAPI for 10 minutes in the dark, and mounted with FluorSave. Images were taken using a Leica DMi8 confocal microscope equipped with a 10× HC PL Apo CS2 objective (NA 0.40) as described before ^46^. Image analysis was performed using the LAS Application Suite.

### Immunofluorescent cell staining

Cell staining experiments were performed as previously described ^46^. MDCK and hCK cells grew on 12mm coverslips until 90% confluency. Cells were fixed with 4% paraformaldehyde in PBS for 20 min at RT and permeabilized with 0.1% Triton in PBS for 5 minutes. HA proteins were precomplexed with α-strep-tag human antibody (GenScript) and goat-α-mouse human antibody (Novus Biologicals) for 30 minutes at a molar ratio of 4:2:1, respectively. For nanoparticle assembly, HA proteins were pooled with CompB in PBS overnight at 4 °C. The protein mixes were then incubated for 90 minutes at room temperature. Cells were washed with PBS, and nuclei were stained with DAPI (Invitrogen) for 10 minutes at RT. Images were taken as described above for tissue staining.

### Quartz Crystal Microbalance with Dissipation monitoring (QCM-D)

SiO_2_-coated quartz sensors (QSX 303, Biolin Scientific) were used for QCM-D measurements. The sensors were cleaned using 2% (w/v) SDS solution by gently heating at 45 °C for 45 minutes, followed by sonication for 5 minutes. The sensors were then washed with copious amounts of water and dried using a nitrogen blow. All clean sensors were activated using 1% Hellmanex solution at 35 °C. QCM-D measurements were performed by using a Q-sense E4 4-channel (Biolin Scientific) with a peristaltic pump (Ismatec IPC ISM930A). The flow rate was adjusted to 30 μL/minute, and the measurement temperature was set to 22 °C.

All solutions described here were prepared using the same batch of PBS buffer for a specific QCM-D run. The stock vesicle solution, 1 mg/mL after extrusion, was diluted to 0.1 mg/mL in 1X PBS buffer, pH 7.4. Supported lipid bilayers (SLBs) were formed by vesicle fusion, in which the vesicle solution was injected onto an activated SiO2 QCM-D sensor. Formation of the SLB after blank PBS rinsing was indicated by a final Δfn change of -24 ± 1 Hz. Next, 0.2 μM SAv was injected, followed by rinsing with blank PBS. Then, 1 μM biotinylated glycan solution was injected, followed by a blank PBS rinse. The nanoparticle assembly and the antibody-precomplexed HA were prepared as described above and injected at 30 μL/minute and 22 °C.

Surface cleaning and activation were the same as those described above for the SiO2-coated quartz sensor with a defined thickness of 50 nm (QSX 335, Biolin Scientific) used in the in-situ QCM-D–SE measurement. QCM-D experiments were performed by using Q-sense Explorer (Biolin Scientific) together with SE measurements that were performed using M-2000 spectroscopic ellipsometer (J.A. Woollam) at a fixed incident angle of 65°. SE data of ψ and φ in the range of 320 – 1000 nm were analyzed using CompleteEASE software (J.A. Woollam). The experimental steps and conditions were kept the same as described above.

## Acknowledgements

This research is funded by the Netherlands Organization for Scientific Research (NWO) through an Open Competition Domain Science M-2 grant (OCENW.M20.106). The authors thank Professor Neil King (Institute for Protein Design, University of Washington, USA), Professor Rogier Sanders, and Kwinten Sliepen (both Amsterdam Medical Center, University of Amsterdam, NL) for supplying the I5350A nanoparticles. The authors thank Professor Nieck E. Benes and Cindy Huiskes (Membrane Science and Technology, MST, University of Twente) for access to and help with the ellipsometry facility, and Maxim Huskens for help with the Python model.

## Data Availability Statement

The data that support the findings of this study are available from the corresponding authors upon reasonable request.

