## Supplemental information for "Self-assembling nanoparticles to assess multivalent interactions between influenza A virus hemagglutinin and glycan surfaces"

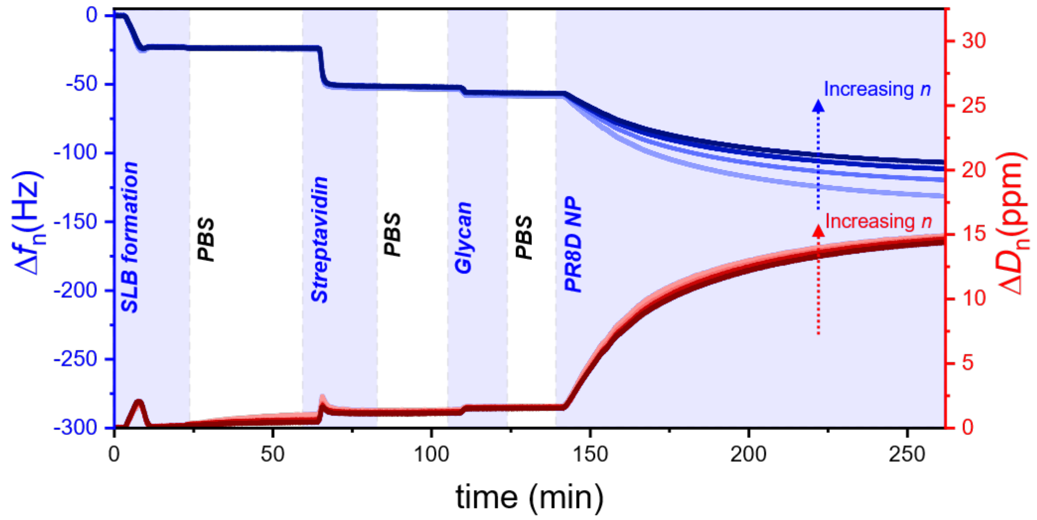


**Figure S1.** An example of QCM-D trace of stepwise formation of SLBs-SAv-glycan layer followed by PR8D190 HA NP binding at 3rd, 5th, 7th, and 9th overtones. The biotinylated lipid fraction was 1.5%, resulting in $\sigma_{SAv}$ of 3.8 pmol/cm$^2$. Glycan density, here of L-6SLN$_3$, is 7.6 pmol/cm$^2$, double the $\sigma_{SAv}$. The concentration of PR8D190 trimer was 10 μg/mL.

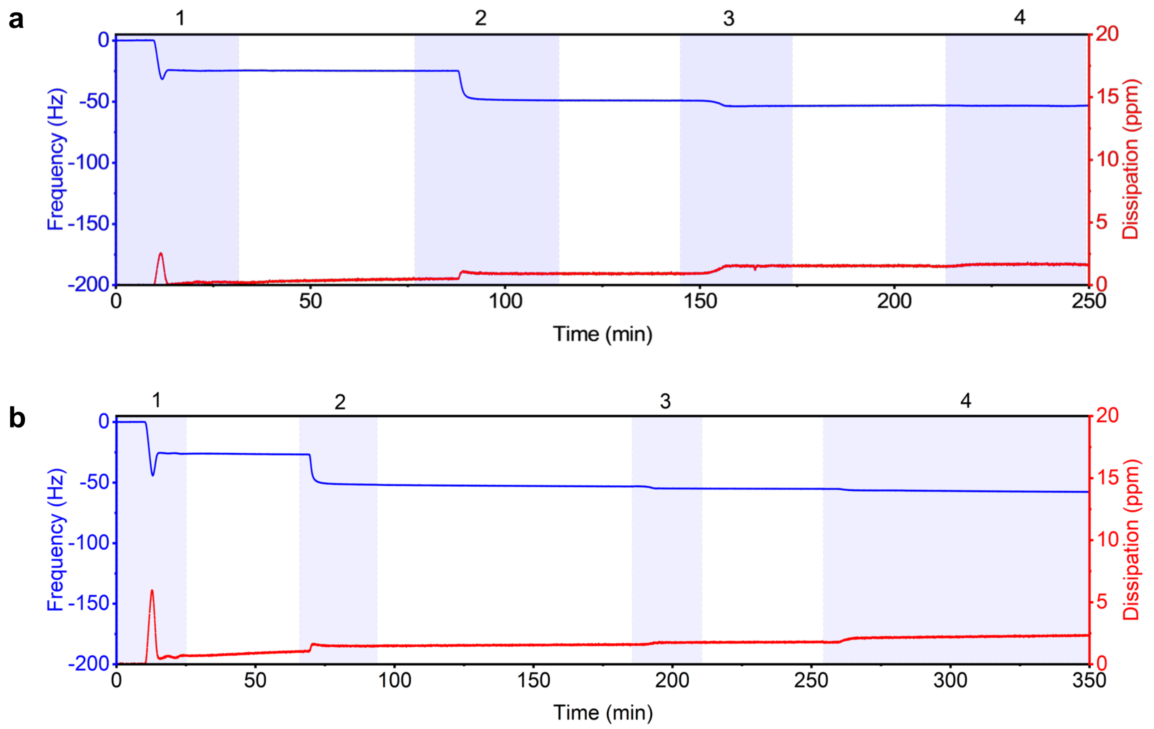


**Figure S2**. SLB formation, glycan immobilization, and binding traces of PR8D190 NP with (a) L-α2,3-SLN3 and (b) L-LN2 (step (4) for both figures). Steps (1), (2), and (3) correspond to the SLB formation, SAv immobilization, and the respective glycan attachment.

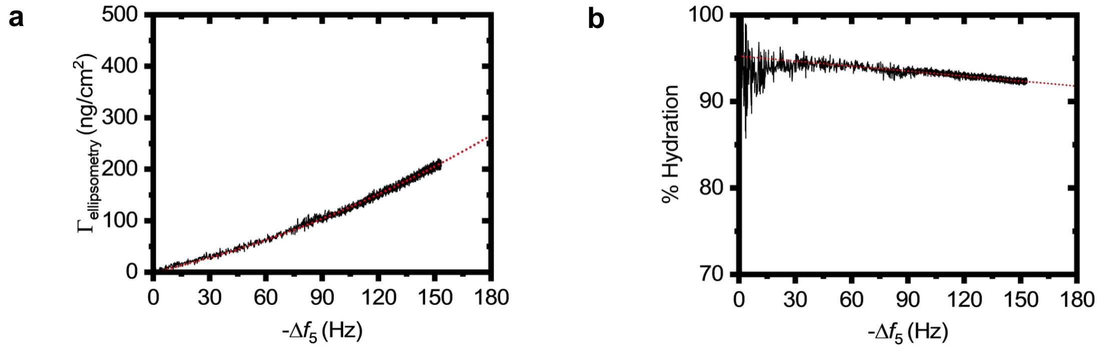


**Figure S3**. (a) Dry mass obtained from spectroscopic ellipsometry (SE) with respect to the QCM response at the 5$^{th}$ overtone. SE and QCM were conducted *in situ*. (b) Percent of hydration (%*H*) as a function of $-\Delta f_5$. %*H* is defined as $\%H = \frac{\Gamma_{QCM} - \Gamma_{ellipsometry}}{\Gamma_{QCM}} \times 100$The linear fit corresponds to the equation of $\%H = -0.0193|\Delta f_5| + 95.26$.

### Estimating the maximum surface coverage of I53-50 HA nanoparticles binding to glycan surfaces

Dry mass information combined with acoustic ratio data from QCM-D enables verification of the expected layer configuration of bound NPs on the surface. The maximum coverage of bound NPs was determined by the $\Delta D_n/-\Delta f_n$ ratio, or the acoustic ratio. For a discrete nanoobject on the surface, the acoustic ratio showed a linear trend and a consistent -Δfn intercept at zero ΔDn/-Δfn across multiple QCM-D overtones.[1–4] This common intercept relates to the close-packing limit of spherical NPs on the surface. The intercept is 263 Hz (**Figure S4**). Converting this frequency shift to dry mass using the relationship shown in Figure S2 results in a dry mass of 461 ng/cm$^2$.

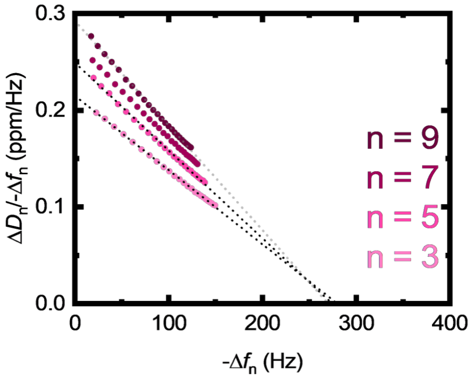


**Figure S4**. Acoustic ratio or the $\Delta D_n/-\Delta f_n$ ratio of PR8D190 NP. We used linear triLacNac α2,6-linked Sia (L-6SLN$_3$) on SLB-SAv layer containing 1.5% biotinylated lipid fraction, resulting in $\sigma_{SAv}$ of 3.8 pmol/cm$^2$. The concentration of the NP was 1 nM. The common intercept across all overtones is 263 Hz.

The nanoparticle is assembled in icosahedral symmetry, resulting in 20 HAs located at the faces, or the HAs sit as the $C_3$ symmetry axes. The nanoparticle is composed of two components, namely N-or PR8D HA triCompA and CompB. One N-or PR8D HA triCompA is composed of trimeric hemagglutinin (80 kDa for each monomer, resulting in a 240 kDa trimer) fused to a mOrange2 (N-terminally) located from hemagglutinin (28 kDa) with a GCN4 (tri, 3 kDa) and CompA (22 kDa). The molecular weight of each N-or PR8D HA triCompA is 396 kDa. CompB is a pentameric protein (90 kDa each), and each nanoparticle has 12 CompB, or can be viewed as the 12 vertices of an icosahedron. Therefore, the molecular weight of each nanoparticle is $MW_{\mathrm{NP}} = (20 \times 396\ \mathrm{kDa}) + (12 \times 90\ \mathrm{kDa}) = 9000\ \mathrm{kDa}\ \mathrm{or}\ 9 \times 10^6\ \mathrm{g/mol}$. Hence, the mass of one nanoparticle can be obtained by dividing the molecular weight by Avogadro's number ($N_A$ = 6.02 × 10$^{23}$ particle/mol), resulting in $m_{\mathrm{NP}} = \frac{9\times10^6\ g/mol}{6.02\times10^{23}\ particle/mol} = 1.5 \times 10^{-17}\ g/particle$.

There is a constraint on the packing of the NP due to the size and exclusion volume. For simplicity, we first assume the upper limit of NP coverage if hexagonal close packing (*hcp*) is adopted. The footprint of the nanoparticle is assumed to be a circle with the same diameter as the nanoparticle itself, $d_{NP} = 2r_{NP}$.

In a *hcp* configuration, the packing area density (of the footprint on a 2D plane, η) can be calculated as written in Equation S1.

$$\eta = \frac{3A_{60^\circ\,\mathrm{sector}}}{A_{\mathrm{equilateral\ triangle}}} = \frac{3\times\frac{1}{6}\pi\left(\frac{d_{\mathrm{HA}}}{2}\right)^2}{\frac{\sqrt{3}}{4}(d_{\mathrm{HA}})^2} = 0.907 \qquad \text{(Eq. S1).}$$

Maximum coverage in *hcp* configuration of the nanoparticle in ng/cm$^2$ is shown in Equation S2.

$$\Gamma_{max.\ hcp} = \frac{m_{NP}}{\frac{1}{\eta}A_{footprint}} = \frac{1.732\times10^6}{d_{NP}^2} \qquad \text{(Eq. S2).}$$

$d_{NP}$ is in nm. The *hcp* configuration should be the upper limit of the possible packing density. Given the dry mass at maximum coverage of the NP is 461 ng/cm$^2$, the calculated $d_{NP}$ using Eq. S2 is 61 nm. This closely matches the diameter of the NP observed in CryoEM in Figure 1B of the main text.

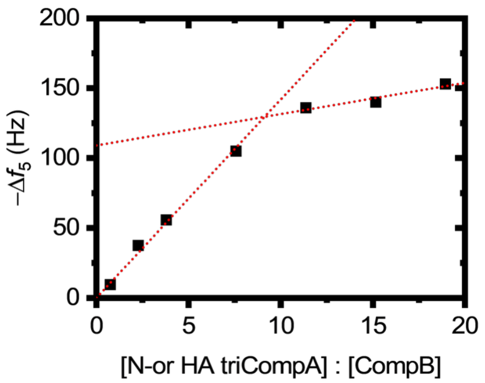


**Figure S5**. -$\Delta f_5$ equilibrium signal with respect to the stoichiometry ratio between trimeric PR8D190 CompA and pentameric CompB. The concentration of CompB was kept constant at 0.46 nM. -$\Delta f_5$ plateaued beyond 7.5-fold PR8D190 CompA. We therefore used this ratio for the rest of the QCM-D experiments.

**Model to describe the multivalent binding of I53-50 HA nanoparticles**

To model the avidity enhancement of multivalent HA presentation with glycans anchored to 2D planar surfaces, we used the model previously described in reference (5). In short, the overall avidity ($K_{av}$) of a multivalent binder can be expressed as written in Equation S3

$$K_{av} = N_A V_{ex} \left(1 + \frac{K_{i,eff}}{N_A V_{explore}}\right)^N \qquad \text{(Eq. S3).}$$

Here, $N_A$ is the Avogadro's number, $V_{ex}$ is the excluded volume of the nanoparticle ($V_{\text{ex}} = \frac{4}{3}\pi r_{\text{NP}}^3$, $r_{NP}$ is the radius of the nanoparticle. $K_{i,eff}$ is the monovalent association constant, $V_{explore}$ is the volume accessible from a single glycan perspective at maximum SAv density, and $N$ is the number of formed bonds dictated by the lowest between the density of binding pockets ($\sigma_L$) or the density of receptors ($\sigma_R$) within the contact area ($A_{contact}$) of the nanoparticle ($N = A_{\text{contact}}\min(\sigma_{\text{L}}, \sigma_{\text{R}})$).

The HA density on the IAV surface was reported to be around 1.3 pmol/cm$^2$ (13 spikes in 40×40 nm$^2$),[6] or around 3.9 pmol/cm$^2$ of available binding sites. In the case of $\sigma_{HA}$ of the IAV, the assumption is that the spacing between binding pockets within an HA trimer is similar to the distance to the binding pocket on the neighboring HA. In other words, the binding pockets are densely packed with no empty gaps in between.

In the case of the I53-50 HA nanoparticle, however, we have to consider the angle between adjacent HAs, enforced by the structure of the assembled NP components. As illustrated in Figure S7a, the HAs are located at the faces of an icosahedron. We assume that the HAs protrude along the normal of each icosahedral face (Figure S7b). The dihedral angle of an icosahedron is 138.2°. The angle between HA (γ) is $\gamma = 180° - \theta_{\text{dihedral}} = 180° - 138.2° = 41.8°$. The end-to-end distance between HAs can be calculated as $d_{\text{HA,NP}} = 2r_{\text{NP}}\sin\left(\frac{\gamma}{2}\right)$. Because of this angle, we also estimate that only 5 HA trimers, located around the apex of a single HA pentamer facet, can simultaneously interact with a glycan surface (Figure 7a, bottom projection). We assume that all HA sites of these 5 trimers are within reach of the glycans present on the surfaces used in this study, while all sites of all other HA trimers are outside this reach. This leads to a total of 15 HA sites that can simultaneously interact with glycans on a 2D surface.

The HA is therefore definitely not as densely packed as in IAVs. By inserting $r_{NP}$ = 31 nm and $\gamma$ = 41.8°, we obtain $d_{HA, NP}$ = 22 nm. The size of an HA head is around 8 nm.[6] The binding pockets are located within the globular head and arranged like an equilateral triangle with a distance of 5 nm. Because $d_{HA, NP}$ is calculated as the HA center-to-center distance, there is a gap of around 14 nm between two HA trimers. This is illustrated in Figure S7c.

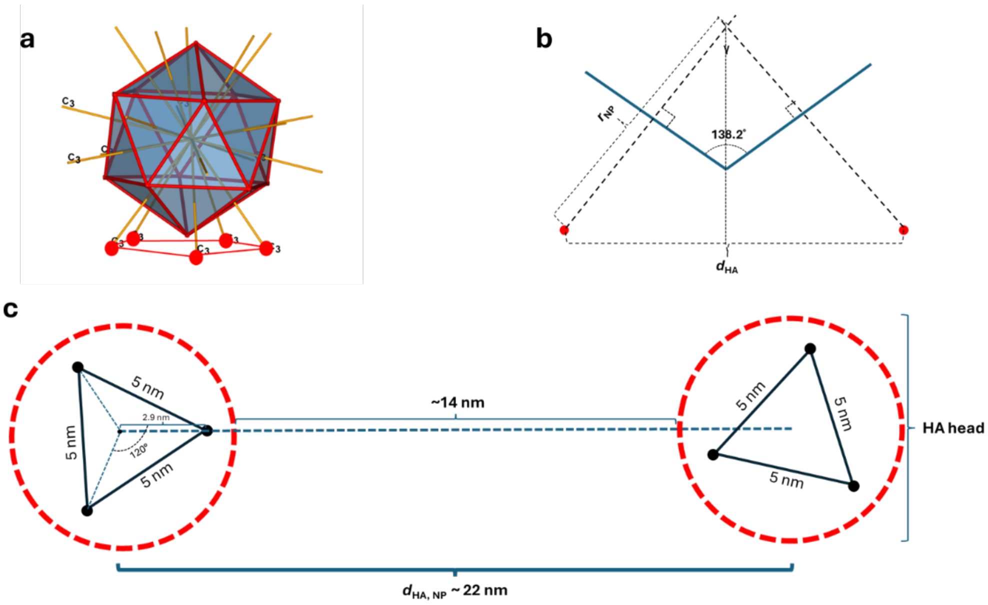


**Figure S6** (a) Representation of an icosahedron facing a surface. (b) Dihedral angle of an icosahedron and geometric representation of the nanoparticle to calculate the distance between two HA heads. (c) Illustration of two HA heads with three binding sites configured in an equilateral triangle with a spacing of 5 nm apart. The binding sites are located within an HA head with the size of 8 nm. Due to the calculated center-to-center HA distance, the HAs are separated with a gap in between, unlike the IAVs.

The surface density of HAs on the I53-50 nanoparticle ($\sigma_{HA,\ NP}$) can be calculated by considering 20 HAs placed on the surface of a circumsphere with a radius of $r_{NP}$. Since there are 20 HAs per nanoparticle, hence. $\sigma_{HA,NP} = \frac{20}{4\pi(31\ \mathrm{nm})^2} \times \frac{1\ \mathrm{mol}}{6.02\times10^{23}} \times \frac{1\ \mathrm{pmol}}{10^{-12}\ \mathrm{mol}} \times \frac{1\ \mathrm{cm}^2}{10^{-14}\ \mathrm{nm}^2} = 0.26$ pmol/cm$^2$ or $\sigma_{\text{binding sites NP}} = 0.78$ pmol/cm$^2$ if one calculates the binding pocket density.

Some implications are: (1) This shows that $\sigma_{HA,\ NP} < \sigma_{HA,\ IAV}$, which implies that the HA nanoparticle binds with less valency compared to IAVs, and the threshold receptor density of the nanoparticle should shift towards larger values than the IAVs. (2) If the receptor density within a contact area of the NP is above 0.78 pmol/cm$^2$, all binding sites within an HA trimer on the nanoparticle can be occupied and thus, $N = A_{contact} \times \min(\sigma_{\text{binding sites NP}}, \sigma_{glycan}) = A_{contact} \times \sigma_{\text{binding sites NP}}$ in Equation S3. (3) We assume the NP binds with a maximum of 5 HAs facing the receptor surface as explained above. The fit results are tabulated in Table S1.

Table S1 Parameters of Eq. S3 used in the fitting of Figure 5A.

| | $K_i$ ($M^{-1}$) | $N_A V_{ex}$ ($dm^3$/mol) | 1 / $N_A V_{explore}$ (M) | $K_i$ / $N_A V_{explore}$ |
|---|---|---|---|---|
| L-6SLN$_3$ | 2.8 × $10^2$ | 7.2 × $10^4$ | 1.4 × $10^{-2}$ | 4.0 |
| N-6SLN$_3$ | | | 2.1 × $10^{-2}$ | 2.9 |

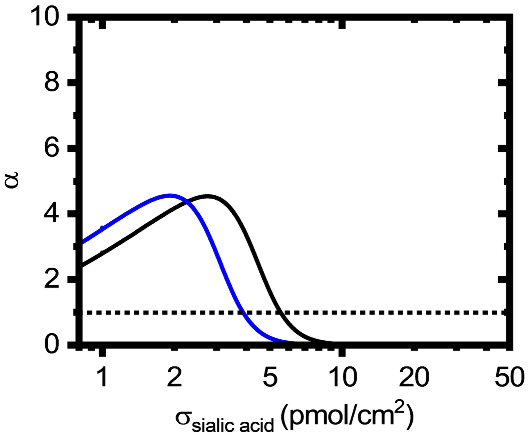


**Figure S7** Selectivity factor (α), defined as $\frac{d\theta}{d\sigma_{\text{sialic acid}}}$ or the slope of log-log plot of coverage-glycan surface density, of L-6SLN$_3$ (black) and N-6SLN$_3$ (blue). The dashed line indicates the α = 1 threshold above which the binding is considered superselective.

**Geometrical model to determine the fractions of I53-50 HA nanoparticles with a minimal binding patch**

For predicting how mixed NPs interact with a glycan surface, we need to estimate which fractions of these particles contain a so-called minimal binding patch, *i.e.,* a patch of 5 HA trimers located around a single apex (see Figure S7), which contains a certain number of functional HAs (ranging from 1-5) that is sufficient to bind to a glycan surface.

I53-50 HA nanoparticles adopt icosahedron symmetry, giving 20 triangular faces, 12 vertices, and 30 edges. Each HA occupies a triangular face, and each vertex forms the apex of a pentamer of triangular faces. There are $2^{20}$ possible nanoparticle combinations of functional/non-functional HAs on the surface of a nanoparticle, of which 17824 are unique due to symmetry. Basic assumptions used here to describe nanoparticle binding on 2D surfaces are: (1) a binding patch is no more than a pentagon consisting of five neighboring triangular faces joined at one shared vertex; (2) particles containing at least one binding patch bind similarly regardless of the number of patches present; (3) on 2D surfaces, a minimum of one binding patch is required for successful binding; for 3D binding at least two such patches need to be present, and these patches have to be located at non-neighboring vertices. Thus, we have to count all particles in all the possible combinations that have at least one binding patch (or two for 3D binding), and do so as a function of the patch size ($P$) of 1, 2, 3, 4, and 5. Binding patches are defined as follows:

- For $P$ = 1: facets with a functional HA,
- For $P$ = 2-5: $P$ functional HAs join at a shared vertex (*i.e.*, within one pentameric binding patch).

For all possible particles (*i.e.*, the $2^{20}$ possible combinations), we have to determine whether it is a binding particle or not. In other words, binding particles should satisfy the requirement above for each $P$. The following steps are taken to evaluate a particle as binding or non-binding:

- Evaluate all 12 vertices per particle
- For each vertex, count all facets with a functional HA
- If the number of functional HAs at a vertex is equal to or greater than $P$, that is a “binding vertex”; if not, it is a non-binding vertex. This yields a list of 12 numbers (either 0 or 1) per particle to indicate the non-binding and binding vertices
- A particle is counted as a “binding particle” for a 2D binding event when it contains at least one binding vertex

- A particle is counted as a "binding particle" for a 3D binding event when it contains at least two binding vertices where these are non-neighboring vertices.

We did this assessment in a Python program that, for each possible particle, determines whether it is binding or non-binding based on the P criteria and whether the binding is 2D or 3D. These particles can be summed to yield the number of binding particles as a function of the total number of functional HAs. Next, we have to calculate the fractions of binding particles as a function of $f_{HA}$ since this fraction of functional HAs is varied experimentally. Thus, we can calculate the fraction of binding particles per *P* as a function of the number of functional HA (between 0 and 20). The probability of forming a particle with a specific number of functional HAs ($Pr_k$) can be calculated by using the binomial probability: $Pr_k = \binom{n}{k} f_{HA}^{k} (1 - f_{HA})^{n-k}$, where *n* is the total number of facets (or HAs, here equal to 20), *k* is the number of facets containing a functional HA (between 0 and 20). The probability of having a functional HA on a facet is equal to $f_{HA}$, which is the experimental fraction of functional HA itself in the mixture. So, for each $f_{HA}$, we need to multiply $Pr_k$ by the fraction of binding particles across all possible combinations with that number of functional HAs, and sum these across all *k*. This yields the fraction of bound particles as a function of fHA for a given patch size *P*. The results for 2D and 3D binding, as a function of *P,* are shown in **Figure S9**.

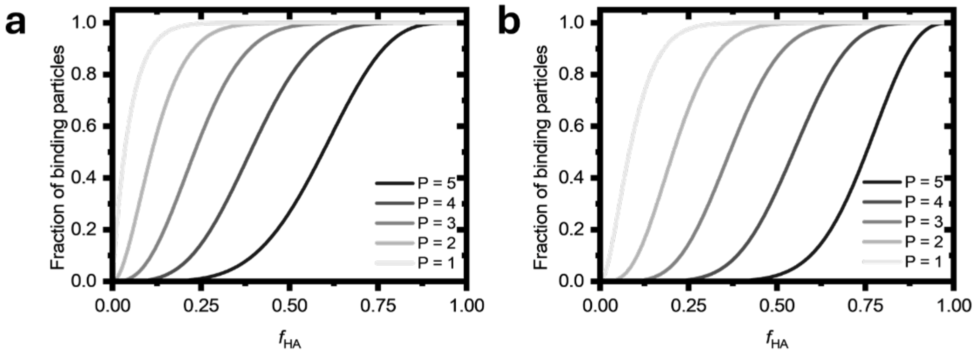


**Figure S8**. Fractions of total particles that can bind in (a) 2D and (b) 3D, as a function of the functional HA fraction and of the minimal binding patch (*i.e.,* the number of functional HAs required to give binding).

For example, the curve for *P* = 5 (meaning the binding requires a minimal binding patch of 5 functional HAs) in **Figure S9a** shows that for $f_{HA} > 0.85$, all particles are "binding", *i.e.*, they have at least one vertex containing 5 functional HAs (i.e., all HA trimers adjacent to that vertex are functional). Lowering $f_{HA}$ to approx. 0.20 reduces the fraction of particles having at least one such binding patch to zero. For *P* = 3 (meaning the binding requires a minimal binding patch of 3 functional HAs), practically all particles are "binding" for $f_{HA} > 0.50$, and the fraction of binding particles drops to zero only when $f_{HA}$ approaches 0. Overall, as the minimum patch size increases (i.e., the HA site density required to achieve binding is higher), a higher fHA is needed to obtain a similar fraction of bound particles compared to a lower patch size.

**Model to describe the minimum number of HA trimers for successful binding of I53-50 HA nanoparticles**

We built a model to describe binding in pure and mixed NP systems. We note that this comes down to assessing which fractions of the particles are binding and which are non-binding, given a certain binding experiment. Here, we varied the fraction of PR8 D190 as the functional HA and PR8 Y98F as the non-functional HA. In the pure NP system, we always have nanoparticles fully decorated with either functional or non-functional HAs, and thus fHA equals the fraction of binding particles. Therefore, the variation of $f_{HA}$ results in a linear concentration variation of the binding NPs, where the concentration of particles that are able to bind, $[NP]_{bind} = f_{HA} * [NP]_{total}$.

For the mixed NPs system, the task is to calculate the fraction of particles in the mixture that can bind at a given $f_{HA}$. These fractions have been determined as a function of the required minimal patch number as described in the previous section. Here, we assume that particles either bind or do not; that is, they bind (with the same affinity) when they contain a minimal binding patch, and they do not bind (zero affinity) without such a patch.

How should we now compare binding results for pure NPs with those of mixed NPs? And can we assess the minimal binding patch for different glycan densities? The trends for binding of pure NPs as a function of $f_{HA}$ are the easiest to read, as we concluded above that these correspond to a linear change in the concentration of binding particles. Thus, these are essentially simple titrations in which the binding particle concentration is varied at a fixed glycan density. We can use these binding responses of the pure NPs to "read" the functional HA fraction of the mixed NPs, thereby achieving the same extent of binding. As explained above, we assume that the fractions of binding particles in the mixed system are determined solely by the fractions of particles with a minimal binding patch, and that their affinities are all equal. Thus, we achieve the same amount of binding when we have the same concentration of binding particles, for both pure and mixed NPs. As an example, we see in **Figure 5C**, bottom-right (17 pmol/cm$^2$), that the pure NPs give half of the maximal binding response at $f_{HA}$ = 0.5. We estimate that we reach the same response for the mixed NPs at approximately $f_{HA}$ = 0.2. Thus, we have a fraction of binding particles of 0.50 at about $f_{HA}$ = 0.2. In **Figure S9a**, this lies on the curve for $P$ = 3. When we perform such an assessment for all data points in **Figure 5C** at all given Sia densities, we obtain **Figure S10**. When we overlay these data points with the fractions of binding particles shown in **Figure S9a**, we estimate that the lowest Sia density (4 pmol/cm$^2$) requires the highest minimal binding patch, most likely $P$ = 5. Similarly, we estimate $P$ = 4 for the intermediate Sia densities and $P$ = 3 for the highest Sia density (15 pmol/cm$^2$).

This confirms that there is a balance between Sia and HA site densities to achieve binding: binding can occur at lower Sia densities only when compensated by higher HA densities, and *vice versa*.

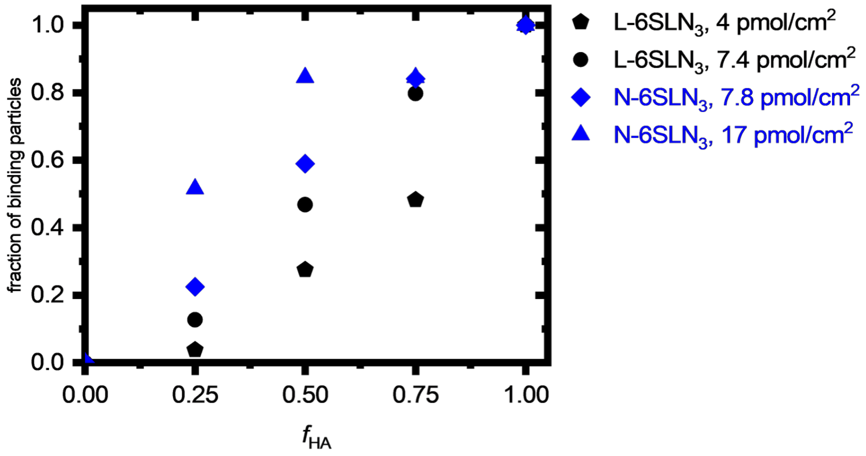


**Figure S10**. Fractions of particles that bind in 2D are calculated from **Figure 5C** for the mixed NP system. Fractions of binding particles are reported as the equivalent concentration of pure, fully functional NPs that yields the same binding response, normalized to the total particle concentration, at fHA = 1 for that particular Sia density. Equivalent concentrations, *C*, are estimated by taking the fraction $f_{HA}$ at which the particle coverage of the mixed system is equal to that of the pure NP system.

Furthermore, mixed NPs bind with an overall lower fraction of binding particles but higher in selectivity (α parameter). Same data points in **Figure 5C** for the mixed NP system are plotted in **Figure S11a** as a function of Sia surface density, combining data points at each $f_{HA}$ to form their own series. Since we conclude that the binding response is the product of both Sia and HA densities, the concentration of binding particles also varies along the superselective curve when fitting the data in **Figure S11a**. As shown, the threshold density shifts to higher Sia density as fHA decreases in both pure and mixed NP systems. A more interesting observation is that the mixed NPs exhibit higher superselectivity than the pure NP system. A reduction in concentration shifts the threshold receptor densities to higher values and changes the quality of superselectivity toward a higher α parameter.[7] As explained previously, the concentration of binding particles decreases in a non-linear fashion in the mixed NP system. On the contrary, it decreases linearly in the pure NP system. In the pure NP system, there is no dependency on the fraction of binding particles, as the receptor density varies, since all particles with functional HAs are binding particles with a maximum patch size of $P$ = 5. However, with the mixed NP system, particles are distributed according to their patch size.

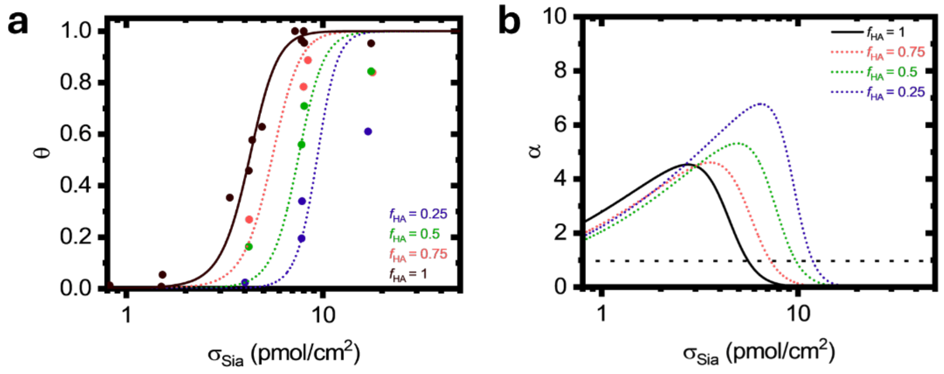


**Figure S11**. (a) Data in **Figure 5C, plotted as a function of the Sia surface density for each fHA series of the** mixed NP system. Dotted lines are the fit using Eq. S3. $f_{HA}$ = 1 is the same data as L-6SLN$_3$ in Figure 5B in the main text. (b) The selectivity parameter from panel (a). The dashed black line is the α value of 1. $f_{HA}$ = 1 is the same as Figure S8.
